# Bacterial filament division dynamics allows rapid post-stress cell proliferation

**DOI:** 10.1101/2020.03.16.993345

**Authors:** Julien Cayron, Annick Dedieu, Christian Lesterlin

**Affiliations:** Microbiologie Moléculaire et Biochimie Structurale (MMSB), Université Lyon 1, CNRS, Inserm, UMR5086, 69007, Lyon, France

**Keywords:** Bacterial filaments, exit from filamentation, cell division, chromosome intracellular organisation, stress response, real-time microscopy in live-cells

## Abstract

Many bacterial species grow into filaments under stress conditions. Initially regarded as an indicator of cell death, filamentation is now proposed to be a transient morphological change that improves bacterial survival in hostile environments. However, the mechanism of filament recovery remains poorly characterized. Using real-time microscopy in live-cells, we analysed the fate of filamentous *Escherichia coli* induced by antibiotic-mediated specific inhibition of cell division, or by UV-induced DNA-damage that additionally perturbs chromosome segregation. Both filament types recover by successive and accelerated rounds of divisions, which are preferentially positioned asymmetrically at the tip of the cell by the Min system. Such division dynamics allows the rapid production of daughter cells with normal size, which DNA content depends on the progression of chromosome segregation prior to division. In most filaments, nucleoid segregation precedes tip-division, which produces nucleated daughter cells that resume normal growth. However, when segregation is deficient, tip-division occurs in the absence of DNA and produces anucleated cells. These findings uncover the mechanism by which bacterial filamentation allows efficient post-stress cell proliferation.

**One Sentence Summary:** Bacterial filaments recover by successive, frequent and Min-dependent asymmetric tip-divisions that rapidly produce multiple daughter cells with normal size and viability

## Introduction

Bacterial proliferation requires coordination between cell cycle fundamental processes, including DNA replication and segregation, cell elongation and division (Donachie, 1968; Helmstetter, 1969; Donachie and Begg, 1970; Donachie et al., 1976; Nicolas et al., 2014; Reyes-Lamothe and Sherratt, 2019). Growth in favorable conditions produces bacterial cells with tightly regulated division rate, size and DNA content (Cooper and Helmstetter, 1968; Campos et al., 2014; Taheri-Araghi et al., 2015; Sauls et al., 2016; Wallden et al., 2016; Willis and Huang, 2017; Micali et al., 2018; Kleckner et al., 2018; Harris and Theriot, 2018; Si et al., 2019). By contrast, unfavorable conditions cause many bacterial species to grow into filamentous cells, formed when division stops while cell elongation continues. Filamentation occurs in response to starvation (Heinrich et al., 2019; Kitahara and Kusaka, 1961; Seeger and Jerez, 1993; Wainwright et al., 1999), salt, pH and temperature shocks (Bereksi et al., 2002; Cameron et al., 2012; Gill et al., 2007; Giotis et al., 2007; de Jong et al., 2004), ionizing radiations (Adler and Hardigree, 1965; Rudolph et al., 2007), or exposure to the host immune response during infection (Justice et al., 2006). Filamentation can also be triggered by exposure to antibiotics (Chatterjee and Raychaudhuri, 1971; Hunt and Pittillo, 1968; Klein and Luginbuhl, 1977; Ryan and Monsey, 1981; Spratt, 1975) or genetic mutations that specifically inhibit fundamental cell cycle processes, such as DNA replication, cell division or DNA recombination (Bi and Lutkenhaus, 1991; Breakefield and Landman, 1973; Ishioka et al., 1998; Mulder and Woldringh, 1989; Van De Putte et al., 1964).

The filamentation phenotype has long been considered as a symptom of stress-induced cell death until the accumulation of evidence that filaments are often able to resume division upon return to favorable conditions (Adler and Hardigree, 1965; Bos et al., 2015; Chen et al., 2018; Dev Kumar et al., 2019; Goormaghtigh and Melderen, 2019; Heinrich et al., 2019; Ishioka et al., 1998; Mulder and Woldringh, 1989; Wehrens et al., 2018; Woldringh and Mulder; Barrett et al., 2019; Miller et al., 2004; Pribis et al., 2019). Filamentation is even part of the normal life cycle of some bacterial organisms, such as *Caulobacter crescentus* in fresh water (Allison et al., 1992; Heinrich et al., 2019), the cyanobacterium *Synechococcus elongatus* growing under dim light (Liao and Rust, 2018) or *Legionella pneumophila* within biofilms (Piao et al., 2006). Two recent studies revealed that *E. coli* filaments (Wehrens et al., 2018), *Synechococcus elongatus* filaments (Liao and Rust, 2018; Wehrens et al., 2018) and *Vibrio parahaemolyicus* (Muraleedharan et al., 2018) divide by asymmetric division regulated by the Min oscillating system, a major regulator of division site positioning in normal cells (Adler et al., 1967; de Boer et al., 1988, 1989; Frazer and Curtiss, 1975; Schaumberg and Kuempel, 1983). These reports suggested a certain level of conservation of the mechanism of filament division in phylogenetically distant Gram-negative bacterial species. Not only filamentation appears reversible, but has also been shown to improve the survival of several bacterial species in specific niches. During infection, filamentous cells perform swarming to evade the immune cells (Allison et al., 1992) and are more resistant to phagocytosis by macrophages in mammalian hosts (Justice et al., 2004, 2006, 2008, 2014). In the environment, filamentous bacteria are also resistant to consumption by protists predators (Hilbi et al., 2007). All together, these findings gradually led to the hypothesis that filamentation could actually be a transient morphological differentiation that improves bacterial survival under hostile conditions. This proposal implicates that filaments have the potential to divide rapidly into a large number of viable cells upon return to favorable growth conditions. However, the mechanism of exit from filamentation and more specifically the fate of daughters cell remain poorly described.

In this work, we use real-time microscopy to analyze the fate of *Escherichia coli* filaments induced by UV-irradiation or exposure to cephalexin antibiotic. UV-irradiation induces DNA damage, mainly pyrimidine dimers that disrupt the progression of DNA replication, thus resulting in the formation of DNA breaks and the induction of the SOS response (Goosen and Moolenaar, 2008; Radman, 1975; Rastogi et al., 2010; Sinha and Häder, 2002; Soubry et al., 2019). The SOS-response in turn induces the production of a range of proteins including SulA (Courcelle et al., 2001; Simmons et al., 2008), which stops cell division by binding to the essential initiator of septum formation, FtsZ (Bi and Lutkenhaus, 1993). Due to the induction and repair of DNA damages, UV-induced filaments exhibit chromosome segregation deficiencies. By contrast, cephalexin antibiotic specifically inhibits cell division by targeting FtsI septum protein (Hedge and Spratt, 1985), but does not affect DNA segregation. We report that these two filament types divide by successive round of divisions, which occur at *one site at the time*, at an accelerated frequency, and preferentially at the tip of the filament, and which rapidly producing multiple daughter cells with normal size and viability. Together with the literature, our results support the view that filamentation reversibility is an efficient process that facilitates post-stress cell proliferation in a variety of bacterial species.

## Results

### Chromosome segregation and cell division in optimal growth conditions

We first characterized the growth of untreated *Escherichia coli* MG1655 cells producing the fluorescently tagged nucleoid associated protein HU-mCherry (Fisher et al., 2013). Time-lapse microscopy performed in fast growth conditions in microfluidic chamber reveals that the nucleoid DNA describes a precise dynamic in the course of the cell cycle (Supplementary video 1 and Figure 1A-1B). Automated cell and nucleoid detection shows that in newborn cells, the chromosome DNA is structured into two nucleoid lobes, which segregate progressively into four lobes as the cell elongate to the division size (Figure S1A-S1B). Localisation of the fluorescently tagged division protein ZapA-GFP (Alexeeva et al., 2010; Goehring et al., 2005; Gueiros-Filho and Losick, 2002) shows that division septa are strictly positioned at midcell, in the DNA-free zone between segregated chromosomes (Figure S1B-S1C). Coordination between chromosome segregation, cell elongation and division allows the symmetric division of the mother cell into two identical daughters cells at regular interval. By monitoring individual cell lineages, we calculated a generation time (τ) of 26.5 min ±4.8 in the microfluidic chamber (Figure 1C), compare to τ = 26.5 min ±6.5 estimated from CFU/ml (colony forming unit) monitoring during exponential growth phase in liquid culture (Figure 1D).

**Figure 1.**
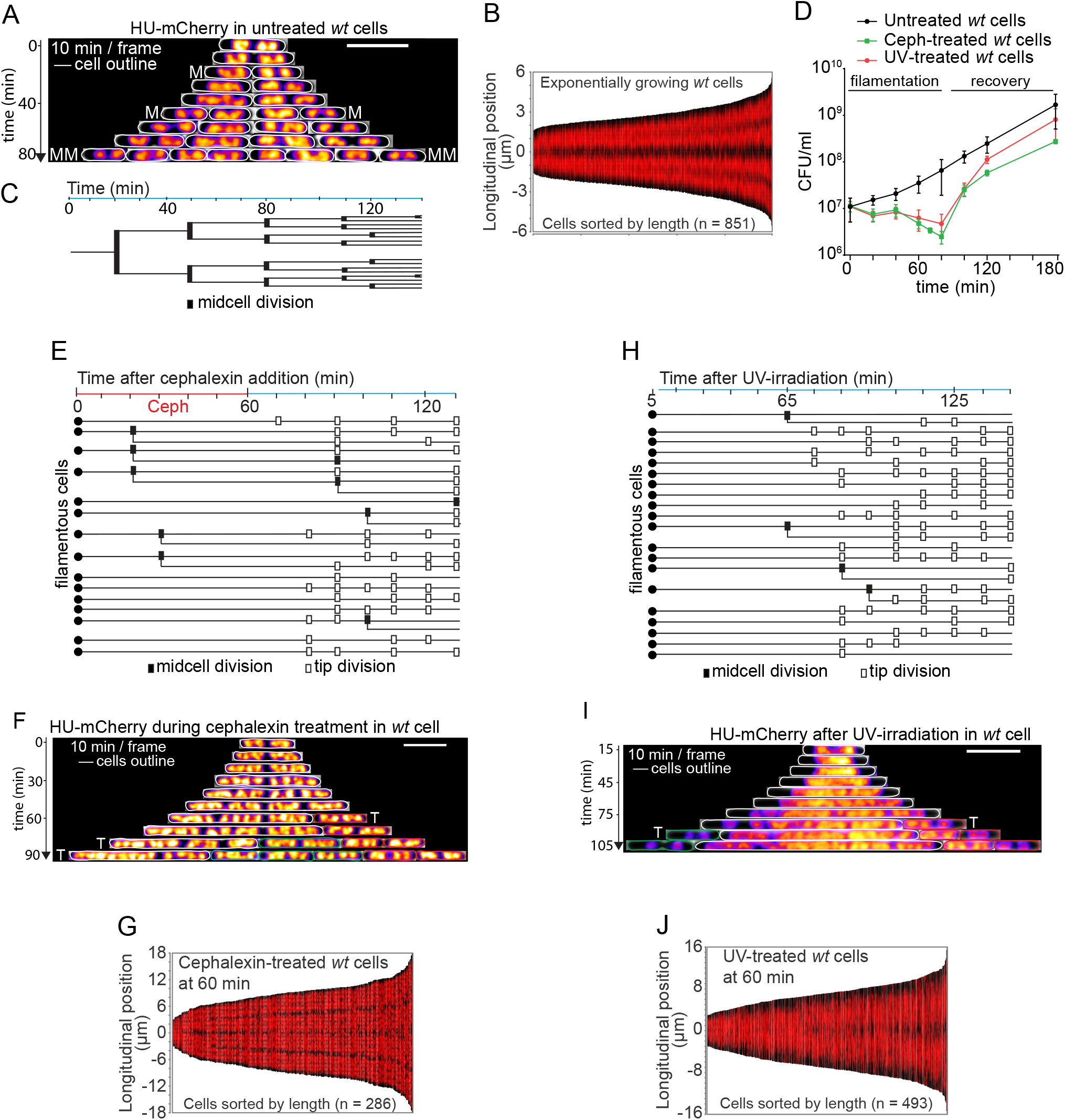
Microscopy analysis the formation and division of UV- and cephalexin-induced filamentous cells. (A) Kymograph of HU-mCherry fluorescence localisation density with the cells outlines (white) of exponentially growing untreated *E. coli* cells visualized by time-lapse microscopy in microfluidic chamber. M indicates midcell divisions on the corresponding frame. Scale bar 5 μm. (B) Demograph of HUm-Cherry localisation in exponentially growing untreated *E. coli* sorted by length. (C) Cell lineage of an untreated *E. coli* cell growing in microfluidic chamber indicating midcell division events. (D) Plot of the CFU/ml over time of untreated (black line), UV-irradiated (red line) and cephalexin-60min-treated (green line) cell cultures. The period of filamentation and recovery are indicated. (E) Timeline representation of division timing and localisation (midcell or tip-division) of individual cephalexin-60min-treated cells monitored by time-lapse microscopy in microfluidic chamber. (F) Kymograph of HU-mCherry localisation density of a representative cell treated with cephalexin for 60 min and then grown in fresh medium. The filament is outlined in white and daughter cells with their lineage are outlined in red and green. T indicate tip-divisions of the cell in the same color code. Scale bar 5 μm. (G) Demograph of HU-mCherry localisation after 60 min of incubation with cephalexin. (H) Timeline representation of division timing and localisation (midcell or tip-division) of individual UV-irradiated cells monitored by time-lapse microscopy in microfluidic chamber. (I) Kymograph of HU-mCherry localisation density of a representative UV-treated cell. The filament is outlined in white. Daughter cells and their lineage are outlined in red and green. T indicates tip-divisions of the cell in the same color code. Time-lapse imaging starts 15 min after UV-irradiation. Scale bar 5 μm. (J) Demograph of HU-mCherry localisation 60 min after UV-irradiation.

### Bacterial population recovery after stress treatment

We then addressed the growth and recovery of *E. coli* populations after UV-irradiation or transient exposure to cephalexin. In our experiments, we use an UV dose (3 J/m^2^) that induces DNA damages provoking 2 logs of viability loss in a RecA-deficient strain, but that are efficiently repaired in a RecA-proficient strain (Figure S2A-S2B). UV treatments trigger the arrest of cell division reflected by the stagnation of the CFU/ml (Figure 1D), associated with the formation of elongated filamentous cells with increased DNA content revealed by time-course microscopy imaging and flow cytometry analysis (Figure S2C-S2E). From ~80 min, the concentration of viable cells increases rapidly (Figure 1D), and cells gradually recover size and DNA content similar to the untreated cells (Figure S2C-S2E). Incubation of *E. coli* cells with cephalexin for 60 minutes results in the similar observations regarding the stagnation and subsequent increase of the concentration of viable cells (Figure 1D), as well as cell morphology and DNA content (Figure S2F-S2H). These two treatments then trigger comparable timing of filament formation and recovery. During the 80 to 120 min time window, we calculated transient doubling times of 9 min ±2.1 for the UV-irradiated populations and 8.2 min ±0.8 for cephalexin-treated cells. This corresponds to a ~3-fold increase of the division rate compare to exponentially growing untreated cells (τ = 26.5 min ±6.5). These results illustrate the reversibility of UV- and cephalexin-induced filamentation, where treated populations eventually recover normal cells length and DNA content. Furthermore, treated populations show a minor reduction of the final concentration of viable cells compare to the untreated population, suggesting that the arrest of division during filamentation is overall compensated by an accelerated production of viable cells during recovery.

### UV- and cephalexin-filaments exhibit different chromosome organisation

We performed live-cell microscopy imaging to get further insight into the process of filament formation and recovery after UV and cephalexin treatments (Supplementary video 1). Single-cell analysis reveals that some cells are able to divide during the first minutes of cephalexin treatment, most probably due to the drug’s inability to inhibit the closure of advanced septa where FtsI proteins (PBP3) are already engaged in cell wall synthesis (Figure 1E) (Eberhardt et al., 2003). These early divisions occur at midcell and produce daughter cells that grow into filaments. Sixty minutes after treatment, the population is mostly composed of filamentous cells harboring regularly separated nucleoids (Figure 1F-1G). By contrast, in UV-irradiated cells the immediate arrest of cell division (Figure 1H) is associated with the disruption of chromosome organisation, reflected by the coalescence of the nucleoids into unstructured DNA masses (Figure 1I and 1J). Within 20 min after irradiation the number of nucleoid detected per cell is reduced to one single DNA mass positioned at the center in ~47 % of the cells, and to two regularly spaced DNA masses in ~42 % of the cells (Figure S3A). These two types of nucleoids conformation after UV-irradiation have been reported previously (Estévez Castro et al., 2018; Odsbu and Skarstad, 2014). By preforming intracellular localisation of the chromosome *terminus*, we show that these outcomes result from the stage of chromosome replication and segregation at the moment of UV treatment. Irradiation of cells with one single *terminus* focus results in cells with one DNA mass, while irradiation of cells with 2 *termini* leads to cells with two DNA masses (Figure S3B-S3D and Supplementary video 2). These results indicate that DNA coalescence is not a global collapse of the whole DNA contained in the cell, but rather occurs intramolecularly within chromosomes that are not fully replicated and separated. Chromosome origin *loci* also tend to cluster together by convergent migration during DNA coalescence (Figure S3E-S3H and Supplementary video 3). Reduction of distances between origins *loci* after UV (Figure S3I-S3J) supports previous proposals that DNA-damage-induced DNA coalescence could correspond to a re-pairing of the replicated sister chromatids to facilitate DNA repair by homologous recombination (Shechter et al., 2013; Vickridge et al., 2017). DNA coalescence is subsequently followed by expansion of the nucleoids into large DNA masses (Figure 1I). Sixty minutes after UV exposure, the population is mainly composed of elongated cells that harbor one or two large unstructured nucleoid DNA structures (Figure 1J). Comparing fate of UV- and cephalexin-induced filamentous cells offers the opportunity to address the influence of nucleoid organisation defects on the process of filament division during recovery.

### Filament division is accelerated and asymmetric

Single-filament lineage analysis reveals several characteristics that are common to the division of cephalexin and UV-filaments. First, filament division occurs *one site at the time* with at most one division per 10 min time-lapse frame interval (Figure 1E and H). The average time lag between two consecutive division events (inter-division time) is 15.8 min ±5.4 for UV-filaments and 14.6 min ±4.9 for cephalexin-filaments with significant heterogeneity, compared to 26.5 min ±4.8 for untreated *wt* cells (Figure 2A). This shows that divisions occur at an accelerated frequently within filaments. Second, analysis of the position of division events reveals that both UV- and cephalexin-filaments preferentially divide asymmetrically at the tip of the filament (referred to as tip-divisions hereafter), and more rarely at midcell (Figure 2B and 2C). Division restart of cephalexin-filaments is relatively synchronous after removal of the drug (Figure 1E and 2B). Tip-divisions of cephalexin-filament always produce nucleated daughter cells that resume normal growth without further forming filaments (Figure 2D and S4A). The midcell divisions observed throughout the recovery process occur between well-separated nucleoids (Figure S4B) and also produced a majority of cells that subsequently divide normally (Figure 2D). Cell division restart is also fairly synchronously in the UV-filaments population (Figure 1H and 2C). Using a plasmid-born *P_sulA_-mCherry* reporter, we show that cell division arrest and restart correlate with the induction of the SOS response, which peaks at 45 min after UV and then decreases to reach basal level at ~75 min (Figure S4C-S4D). The termination of SOS induction further suggests that most SOS-inducing RecA intermediate associated with DNA damages are resolved in that time window. Consistently, we observe that UV-filament division restart is concomitant with the gradual reorganization of individual nucleoids that separates from the edges of the unstructured DNA masses (Supplementary video 1, Figure 1I and S4E). Most tip-divisions consequently produce nucleated cells that resume normal growth (Figure 2D). However, we also observe a significant proportion of anucleated cells produced when tip-divisions occur in DNA-free polar regions of the filament resulting form DNA segregation deficiencies (Supplementary video 1, Figure 2D and S4E-S4F). Besides, the minority of division that occur at midcell are observed in cells where DNA coalescence led to two DNA bulks separated by a central DNA-free space (Figure S4F-S4G). These divisions mostly generate cells that later divide normally, but also frequently produce cells that further form filaments (Figure 2D). Noteworthy, we do not observe the complete resolution of UV-filaments over the duration of our experiments (up to 215 min). Rather, we observe that multinucleated filaments keep growing and dividing sequentially, at accelerated frequency and preferentially from the tip, serving as a mother cell that produce several generations of daughter cells (Supplementary video 1).

**Figure 2.**
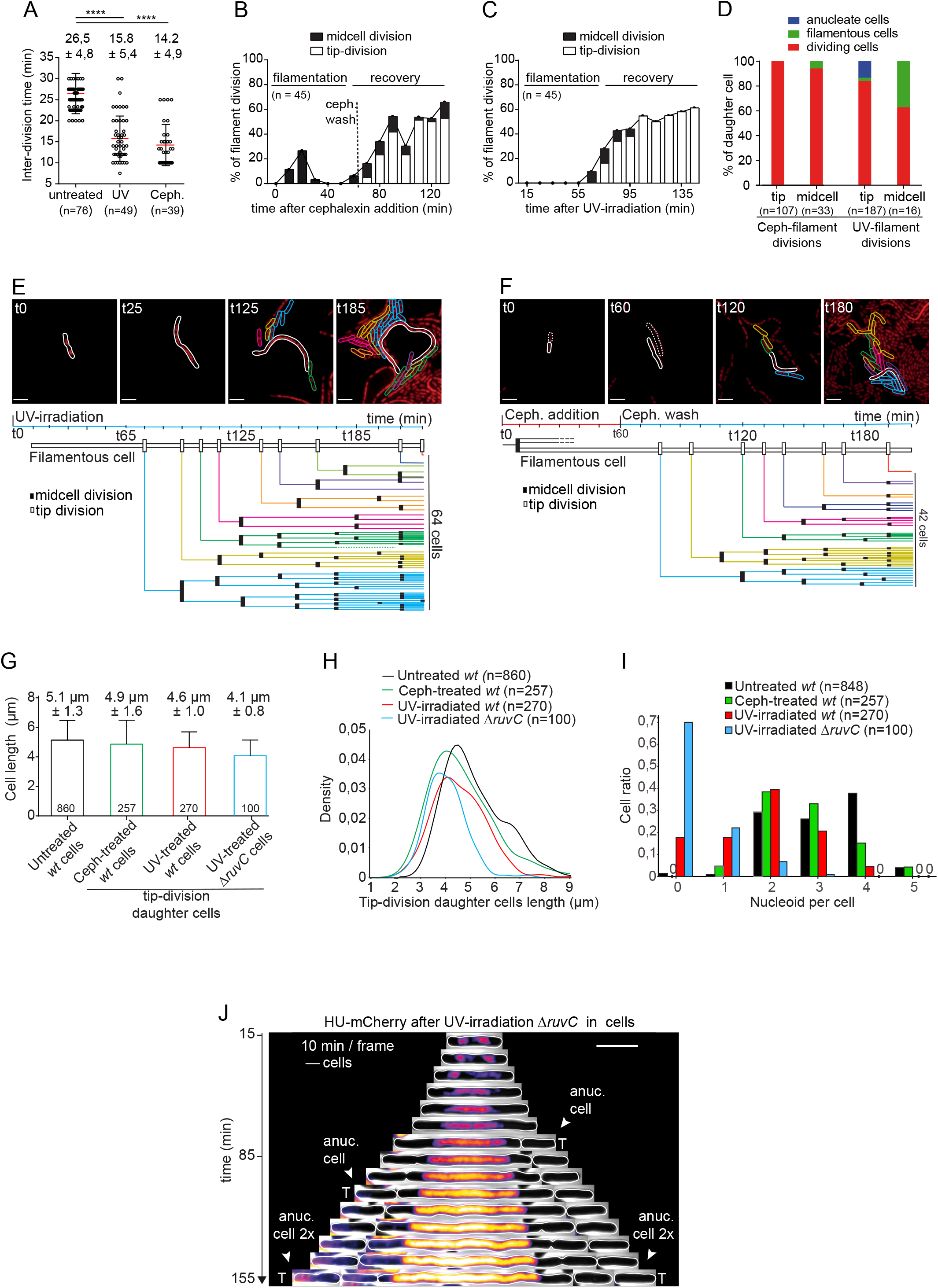
Filaments mainly divide asymmetrically from the tip and produce viable daughters cells with normal length. (A) Time lag between consecutive divisions (inter-division time) in unstressed wt cells, cephalexin-filaments and UV-induced filaments. Mean (red) and standard deviations (black) on (n =) divisions events analysed are shown and indicated above. P-Value significance from unpaired statistical t-test is indicated (**** <0.0001 significant). (B) and (C) Quantification of the percentage of cells dividing (black line) in the course of time-lapse imaging (10 min/frame) during 60 min incubation with cephalexin (B) or after UV-irradiation (C). The split histogram indicates the proportion of midcell (black bar) or tip-division (white bar) of the filament (n = filaments analysed). (D) Histograms presenting the categories of daughter cells resulting from midcell or tip-divisions of the filaments over the entire duration of the experiment (145 min). Anucleated cells are daughter cells lacking HU-mCherry labelled nucleoids. Filamentous cells are daughter cells that in turn grow into filaments. Dividing cells are daughter cells that resume normal division without forming further filaments. (n= cells analyses). (E) Cell lineage of a representative UV-induced filament indicating the timing and location of division events. The lineage of the resulting daughters cells are indicated in distinct colors as well as the total number of cells eventually produced. (F) Cell lineage of a representative cephalexin-induced filament. (G) Average cell length and (H) cell length distribution of exponentially growing untreated *wt* cells (black), daughter cells produced by tip-divisions of *wt* cephalexin-filaments (green), and of *wt* (red) Δ*ruvC* (blue) UV-induced filaments. Error bars indicate the standard deviations. The number of cells analysed is indicated within the corresponding bars. (I) Histogram of the number of automatically detected nucleoid in exponentially growing untreated *wt* cells (black), daughter cells produced by tip-divisions of *wt* cephalexin-filaments (green), and divisions of *wt* (red) Δ*ruvC* (blue) UV-induced filaments. (J) Kymograph of HUm-Cherry fluorescence localisation density with the cells outlines (white) of a representative Δ*ruvC* visualized by time-lapse microscopy in microfluidic chamber. T indicates tip-divisions on the corresponding frame. Anuc. cell indicate the formation of anucleated cell. Scale bar 5 μm.

### Filament division rapidly produces multiple cells with normal length and viability

We next addressed the fate of daughter cells by analyzing the lineage of individual filaments during recovery. Population analysis presented above reveals a certain level of heterogeneity in the dynamics of filament divisions, we then present results for representative individual UV- and cephalexin-filaments. The UV-filament tip-divisions start at 75 min and subsequently occur every 15.5 min on average, resulting in the production of a total of 9 daughter cells (Figure 2E and Supplementary video 4). All these daughter cells subsequently divide normally from midcell and eventually generate a total of 64 cells. This corresponds to an overall generation time of 26.9 min over the duration of the experiment. Similarly, cephalexin-filament tip-divisions start at 80 min and occur on average every 15 min afterwards (Figure 2F and Supplementary video 5). The 8 resulting daughter cells divide from midcell into 42 cells, corresponding to a generation time of 30.9 min over the duration of the experiment. This single-cell analysis confirms that filaments repeatedly divide from the tip to generate daughter cells that further exhibit normal growth dynamics. Furthermore, we observed that tip-division generate daughter cells with length distribution and average remarkably similar to untreated cells (Figure 2G-2H). This strongly suggests that tip-division positioning is regulated by a specific mechanism that controls daughter cells size.

### Filament tip-division events occurs independently of nucleoid organisation

We run automated detection to quantify the number of nucleoids per daughter cells resulting from the filament tip-divisions (Figure 2I). In all cephalexin-filaments and most UV-filaments, nucleoid segregation precedes tip-division. However, the resulting progeny of newborn daughter cells contains a variable number of nucleoids (Figure 2I), which intracellular nucleoid positioning is imprecise (Figure S5A). Therefore, tip-division daughter cells are dissimilar to unstressed newborn cells, which contain 2 well-positioned nucleoids. This indicates that the regulations at play during tip-divisions exert a tight control on the size of daughter cells, but a rather lose control on their nucleoid content and organisation. This interpretation is further supported by the production of anucleated daughter cells when nucleoid segregation is defective prior to tip-division, as in a significant proportion of the UV-filaments that exhibit DNA-free poles (Figure 2D, 2E and S4E-S4F). To exacerbate and further investigate the anucleated division phenotype, we used Δ*ruvC* cells in which the resolution of recombination intermediates is impaired (Ishioka et al., 1998; Otsuji et al., 1974). UV-irradiation of Δ*ruvC* cells leads to the formation of filaments exhibiting large DNA-free tips, due to the inability to repair and segregate the damaged chromosomes. Strikingly, these filaments divide by successive rounds of tip-divisions, all of which occur in DNA-free regions and fail to entrap DNA in the daughter cells, which are all anucleated (Figure 2I-2J and Supplementary video 6). Noteworthy, the resulting anucleated daughter cell population is enriched in smaller cells and exhibits a reduced average size compared to nucleated daughter cells (Figure 2G-2H). These observations are consistent with the view that tip-divisions positioning and licensing do not require the presence of nucleoid DNA at the site of division. Yet, when DNA is present at the site of septum constriction, septum closure is slightly delayed and daughter cells larger.

### Septum positioning in filaments and regulation by SlmA and MinCDE systems

To get further insight into the regulation of the filaments division, we analyzed the positioning of division rings by visualising the septum protein ZapA-GFP. UV- and cephalexin-filaments exhibit a number of ZapA-rings per cell that increases with the cell length (Figure 3A-3B). Most ZapA rings are positioned close to the cell pole. We calculated that 78 % of division events occur at these polar ZapA-rings in UV-filaments, compared to 47.4 % in cephalexin-filaments where midcell divisions are more frequent. Importantly, the distance between polar ZapA-rings that eventually lead to division events is maintained regardless of the filament length (4.0 μm ±1.2 in UV-filaments and 4.6 μm ±1.2 in cephalexin-filaments) (Figure 3C) and is reminiscent of the size of the resulting daughters cells (Figure 2G).

**Figure 3.**
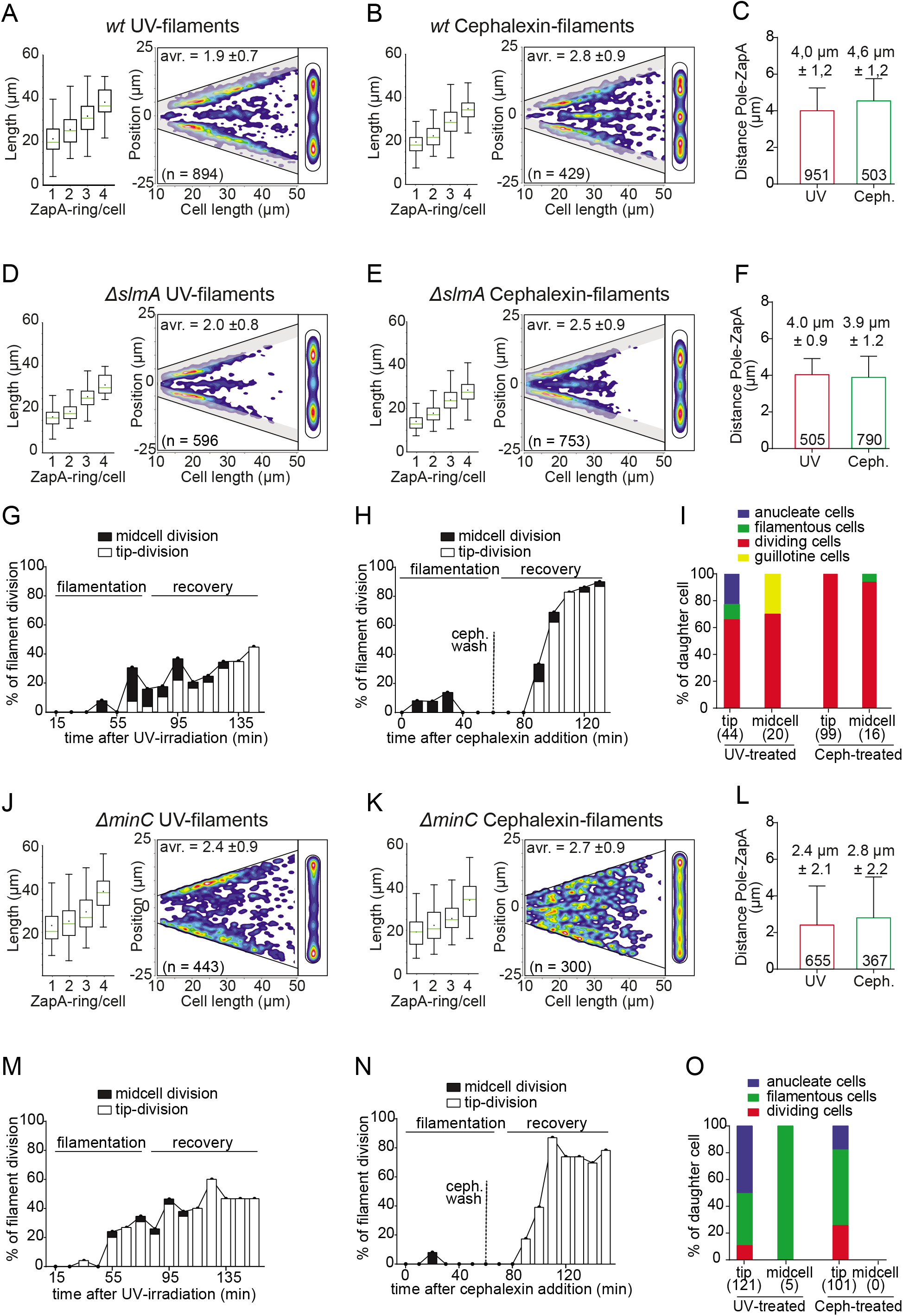
Role of SlmA or MinC on the regulation of filament divisions. (A), (B), (D), (E), (J), and (K) panel present the average cell length of cell with 1 to 4 ZapA-rings (left), ZapA-rings localisation density plot in filaments sorted by length (middle), and ZapA heat map in normalized cell (right). Results are shown for UV-induced filaments (A, D and J) and cephalexin-induced filaments (B, E, and K) in *wt* (A and B), Δ*slmA* (C and D) and Δ*minC* (J and K) strains. The strains and treatments are indicated within the panel with the number of cells analysed (n =). (C) (F) and (L) average distance between polar ZapA-rings and the cell pole in UV-(red) and cephalexin-induced (green) filaments. The error bars indicate the standard deviation on the number of cells indicated within the bars. (G), (H), (M), (N) Quantification of the percentage of cells dividing (black line) in the course of time-lapse imaging (10 min / frame). The split histogram indicates the proportion of division occurring at midcell (black bar) or at the tip of the filament (white bar) (n = 25 filaments analysed for each plot). (I) and (O) Histograms presenting the categories of daughter cells resulting from midcell or tip-divisions of the filaments. Blue, green and red categories are as in to figure 2B, with the addition guillotine cells (yellow) resulting from closing of the septum on the nucleoid DNA cells. (n= cells analyses).

We then addressed the influence of SlmA protein that mediates nucleoid occlusion mechanism (Bernhardt and de Boer, 2005). ZapA-rings are still preferentially located asymmetrically, with comparable distance to the filament pole (Figure 3D-3F). The most significant effect of SlmA inactivation is to increases the rate of unproductive divisions of UV-filaments. Tip-divisions generate more anucleated and filamentous cells and midcell divisions often lead to guillotining of the DNA by the septum (Figure 3G, S6A-6B and Supplementary video 7). The reduced proportion of large filaments also supports that divisions are more frequently licensed in the Δ*slmA* filaments (Figure 3D-3E). These results indicate that SlmA has a minor role in septum positioning within the filament, but acts to avoid septum closing on the unsegregated DNA. Consistently, SlmA inactivation has little impact on the productivity of cephalexin-filament division where chromosomes are regularly separated (Figure 3H-3I) and on the recovery of normal cell size after stress treatments (Figure S6C-6D). By contrast, inactivation of the Min system in Δ*minC* strain dramatically disturbs ZapA localisation and the outcome of filament division. In both UV- and cephalexin-filaments, most ZapA rings are located abnormally close to the cell pole (Figure 3J-3L), leading to mostly polar divisions (Figure 3M-3N) that generate anucleated minicells or cells that further form filaments (Figure 3O and S6E-S6F). Additional ZapA rings positioned along the filament are observed in Δ*minC* cephalexin-filaments and not UV-filaments, most probable reflecting the influence of nucleoid occlusion. These results show that the Min system is a key regulator of septum positioning within filamentous cells. Min-dependent positioning of division events at the tips is critical to the recovery of normal cell size and growth after stress treatment.

## Conclusion

Our results show that cell filamentation is a reversible morphological change that renders cell division transiently dispensable, while allowing efficient post-stress bacterial proliferation (Figure 4). Filamentous *E. coli* divide following a precise sequence of events that primarily involves successive and frequent rounds of divisions preferentially located asymmetrically at the tip of the filament. This division dynamics allows the rapid production of a large number of viable daughters cells that resume normal growth dynamics. Evidence for tip-division of bacterial filaments has been previously reported in *E. coli* (Adler and Hardigree, 1965; Mulder and Woldringh, 1989; Wehrens et al., 2018), *Synechococcus elongatus* (Liao and Rust, 2018) and *Vibrio parahaemolyicus* (Muraleedharan et al., 2018). To address the potential conservation of this mechanism in other Gram-negative bacteria, we investigated cephalexin-induced filaments of both *Salmonella enterica* Thyphimurium LT2 and *Citrobacter rodentium* ICC168 strains and showed that they also appear to divide asymmetrically from the tip of the filament (Figure S7A-S7C). Together with other reports, these observations support that asymmetric tip-division could be a conserved mechanism of filament recovery in various bacterial species.

**Figure 4.**
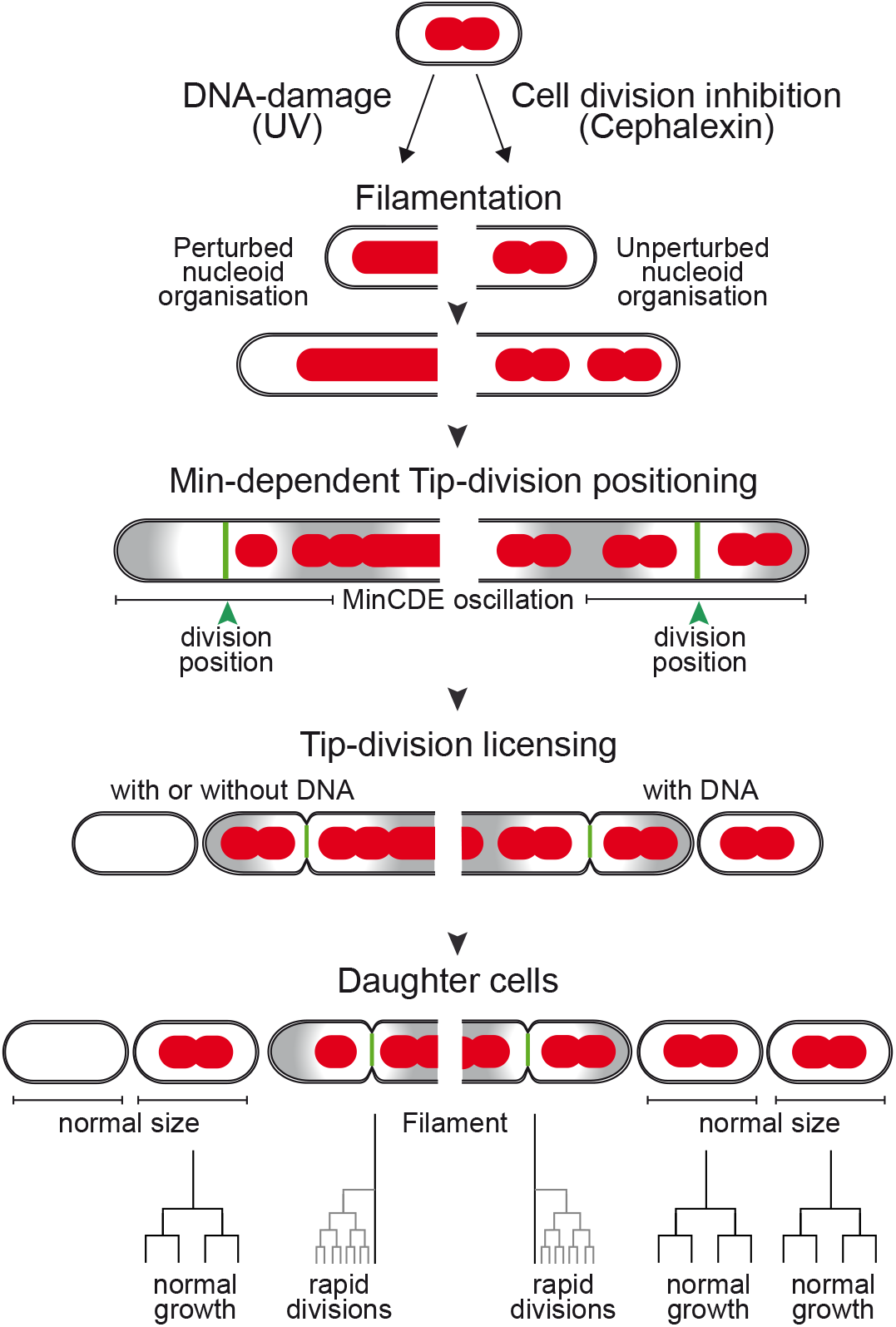
Summary model for division of filamentous cells. UV-irradiation and cephalexin treatments trigger the formation of filamentous cells, which exhibit perturbed and unperturbed nucleoid organisation, respectively. In both filament types, the Min system regulates the positioning of the division site at a distance from the cell pole corresponding to half the oscillation wavelength of the MinCDE system. Tip-divisions occur one at the time, regardless of the presence of nucleoid DNA and lead to the production of daughter cells of normal size. Nucleated daughter cells subsequently exhibit normal growth and viability.

In *E. coli*, asymmetric division has been reported in filaments induced by DNA-damage induction using UV (Rudolph et al., 2007) or exposure to antibiotics such as ciprofloxacin (Bos et al., 2015) and ofloxacin (Barrett et al., 2019; Goormaghtigh and Melderen, 2019). Our results show that tip-division is still the preferential mode of division of DNA damage-free filaments exhibiting well-segregated nucleoids. This excludes the possibility that tip-divisions simply result from the occlusion of midcell division by unsegregated damaged chromosomes. This is also supported by the observation that SlmA inactivation has little effect on the position of division events within UV-filaments. By contrast, we show that the Min system *per se* is key to regulate the positioning of the division sites at the tip of both UV- and cephalexin-filaments. This is consistent with previous study reporting the role of the Min system in the division of filamentous forms of *E. coli* (Wehrens et al., 2018), *S. elongatus* (Liao and Rust, 2018), and *V. parahaemolyicus* (Muraleedharan et al., 2018). In filamentous *E. coli*, MinC oscillation wavelength was estimated between 8 and 10 μm (Ramm et al., 2019; Touhami et al., 2006), meaning that Min minimums are expected to locate 4 to 5 μm from the cells poles. This is consistent with the positioning of polar ZapA-rings and the size of tip-division daughter cells in both UV and cephalexin conditions. Besides, we also show that most filaments harbor multiple ZapA-rings, but division is preferentially licensed at ZapA-rings that are closest to the poles. The reason for division licensing at polar sites remains unclear, yet one potential explanation could lie in the dynamics of Min oscillations within filaments. In the course of the filament elongation, Min gradient positions are constantly readjusted as respect to the cell poles that act as the oscillation barrier (Wehrens et al., 2018). Min gradients are then expected to be more stable at the cell poles than along the filament, which could potentially facilitate the establishment of the division machineries and subsequent licensing. We also establish that tip-division positioning and licensing occur similarly regardless of the presence or the absence of nucleoid DNA at the division site, resulting in the production of daughter cells with tightly controlled normal size. In most filaments, nucleoid segregation precedes tip-division, resulting in the production of nucleated daughter cells that resume normal growth. Yet, when segregation is deficient prior to division, division occurs at DNA-free tip of the filament and fails to entrap DNA in the daughter cell, which is then anucleated. This reveals that division and chromosome segregation are concomitant processes, which can be spatially uncoupled within filaments. Yet, these two processes or not strictly independent, since our data indicates that the presence of nucleoid DNA at the division site negatively regulates septum closure, which is consequently delayed.

If our work uncovers the mechanism of exit from filamentation, other reports establish that filamentation *per se* improves bacterial survival during infection or in the environment (Allison et al., 1992; Hilbi et al., 2007; Justice et al., 2006, 2008, 2014). In addition, filamentation has also been shown to facilitate bacterial evolution and adaptation under stress. Indeed, stress conditions increase the mutation rate within the filamentous cells, thus resulting in a progeny of cells the inherited genetic mutations. This genetic variability was shown to enhance the resistance of the daughter cells to antibiotic treatment (Bos et al., 2015; Pribis et al., 2019). Evidence for filamentation efficient recovery, direct selective benefits, and conservation supports the view that this morphological differentiation plays an important role in the survival and proliferation of various bacterial species within hostile environments.

## Acknowledgments

The authors thank the National BioResource Project, Coli Genetic Stock Center, F. Cornet, Y. Yamaichi, E. Gueguen and E. Cascales for providing strains; Sarah Bigot for critical reading of the manuscript, and A. Ducret for valuable help with MicrobeJ and for writing custom analysis codes.

## Author contributions

C.L. and J.C. conceived the project, designed the study, analysed the data and wrote the paper; C.L., J.C. and A.D. performed the experiments and made the bacterial strains and plasmids.

## Methods

### Bacterial strains, plasmids and growth conditions

#### Strains construction

Strains and plasmids used in the study are listed in Table S1. Fusion of genes with fluorescent tags and gene deletion were constructed using lambda Red recombination method (Datsenko and Wanner, 2000). Chromosomal gene *loci* were transferred by phage P1 transduction to the wanted background strains. When required the *kan* and *cat* resistance genes flanked by FRT sites were removed by expression of the Flp recombinase from plasmid pCP20. Unless otherwise stated cells were grown at 37°C in LB agar medium (Tryptone 10 g/L, Yeast extract 5 g/L, NaCl 10 g/L and Agarose 15 g/L) or grown in MOPS EZ Rich Defined Medium, call RDM (Growth rich medium, 10x MOPS Mixture, 0,132 M K2HPO4, 10x AGCU, 5x Supplement EZ, 20% Glucose, Filtered at 0,22 μm, Teknova). When appropriate, antibiotics were used at the following concentrations: Ampicillin (Ap) 100 μg/mL, Chloramphenicol (Cm) 20 μg/mL, Kanamycin (Kn) 50 μg/mL, Streptomycin (St) 20 μg/mL, Cephalexin (Ceph.) 5 μg/mL.

#### Induction of cell filamentation

Induction of cell filamentation was performed on culture grown in RDM at 37°C at final OD_600nm_ of 0.2 by UV-irradiation or by cephalexin treatment during 60 minutes. For UV-irradiation, the cell culture was transferred into a petri dish and irradiated at 3 J/m^2^ using the UV-CROSS-LINKER CL-508 (Uvitec Cambrige). After UV-irradiation, culture was transferred into clean tube for further growth and sampling. Cephalexin treatment was induced by addition of cephalexin at final concentration of 5 μg/mL during 60 minutes. Cephalexin was then washed away by pelleting cells in 15 mL tubes using centrifugation (2500 rpm 5 min), and the cell pellet was resuspended in an equal volume of fresh medium by gentle pipetting. Cells were then transferred into clean tube for further growth and sampling.

### Spot assay and CFU experiment

Cells were grown overnight in LB at 37°C and serial diluted. 10 μL drop of each dilution was deposited on LB agar plates. The viability of *wt and recA-* strains after UV-irradiation was estimated by comparing concentration of colony forming unit (CFU/ml) on the plates that were or were not irradiated using UV-CROSS-LINKER CL-508 (Uvitec Cambrige) and incubated overnight at 37°C. For monitoring of the CFU/ml after stress treatments (Figure 1D), overnight cultures were diluted to an OD_600nm_ of 0.01 and grown with shaking at 37°C to a final OD_600nm_ of 0.2. Cell culture was treated like described in the *Induction of cell filamentation* section. Samples were collected at the indicated times, serially diluted and spotted on an LB agar plate. Plates were incubated overnight at 37°C and CFU/ml were determined. For the 3 conditions tested (untreated, UV-irradiated and cephalexin-60min-treated), samples corresponding to t_0min_ were normalized at 10^7^ CFU/mL. In order to correct for the loss of cells during centrifugation and resuspention, the CFU/ml after cephalexin wash were normalized to the CFU/ml at 60 min before cephalexin wash.

### Live-cell microscopy imaging and analysis

#### Cell imaging on agarose-pad

Overnight culture of cells were diluted to an OD_600nm_ of 0.01 and grown with shaking at 37°C to an OD_600nm_ of 0.2. Treated or untreated cells cultures were collected at time points indicated and 10 μL samples were deposited on 1% agarose-RDM pad (Lesterlin and Duabrry, 2016) and imaged by microscopy snapshot or time-lapse with 2.5 min/frame. For UV treatment, time-lapse acquisition starts 2.5 min after irradiation.

#### Time-lapse in microfluidic chamber

Time-lapse in microfluidic chambers were performed as described previously (Nolivos et al., 2019). Here, overnight cultures of cells were diluted to an OD_600nm_ of 0.01 and grown at 37°C to an OD_600nm_ of 0.2. UV-treated or untreated cells were immediately loaded into a B04A microfluidic chamber (ONIX, CellASIC®) preheated at 37°C. Nutrient supply was maintained at 1 psi over the duration of the time-lapse imaging (~4 hours). For time-lapse in the presence of cephalexin, cephalexin was injected into the microfluidic chamber during 10 min at 2 psi, followed 50 min at 1 psi. Cephalexin was then washed out by injection of fresh RDM medium during 10 min at 2 psi, followed by 1 psi during 4 hours. For all timelapse in microfluidic chambers, cells were imaged during 4 hours at 10 min/frame. For UV treatment, time-lapse acquisition starts 15 min after irradiation.

#### Image acquisition

Conventional wide-field fluorescence microscopy imaging was carried out on an Eclipse Ti-E microscope (Nikon), equipped with x100/1.45 oil Plan Apo Lambda phase objective, FLash4 V2 CMOS camera (Hamamatsu), and using NIS software for image acquisition. Acquisition settings were 200 ms for sfGFP, 100 ms and 300 ms for mCh for snapshots and time-lapses on agarose-pad and time-lapse in microfluidic chamber, using 50% power of a Fluo LED Spectra X light source at 488 nm and 560 nm excitation wavelengths.

#### Image analysis

Microscopy images processing was performed using the open source ImageJ/Fiji (download at https://fiji.sc/) and quantitative image analysis was performed using the free MicrobeJ plugin (download at http://microbej.com)(Ducret et al., 2016). Cells outlines were detected automatically based on the segmentation of phase contrast image. When required, cell outlines were corrected using the Manual editing interface of MicrobeJ plugin. Cell size was automatically extracted and plotted on GraphPad Prism software. Nucleoids were detected automatically using the Feature detection interface of MicrobeJ plugin (tolerance = 500) and the nucleoids number and area were extracted and plotted over the time on GraphPad Prism software. ZapA-GFP and ParB-GFP (*ori* and *ter*) localisation was performed by automatic detection using the Feature detection interface of MicrobeJ plugin (tolerance = 60 for ParB^P1^-GFP and 500 for ZapA-GFP). Localization of ZapA-GFP was automatically extracted and density plots and heat-map plots were generated with MicrobeJ plugin. Distance between ZapA-GFP rings and the cell pole was extracted automatically and plotted on GraphPad Prism software. Demographs and Kymographs of mCherry fluorescence distribution were generated automatically using MicrobeJ. For analysis of *P_sulA_-mCherry* intracellular intensity after UV was extracted from MicrobeJ and the fold-increase of fluorescence signal was plotted over time on GraphPad Prism software. Analysis of the cell lineages and identification of the type of filament divisions (midcell and tip-division) were performed manually by following individual cells on time-lapse microscope images.

### Flow cytometry analysis

Overnight cultures were diluted to an OD_600nm_ of 0.01 and grown with shaking at 37°C to an OD_600nm_ of 0.2. Samples of treated and untreated cultures were collected at time points indicated and diluted to a concentration of 30000 cells/μL (corresponding to an OD_600nm_ ~0.06) in fresh medium at 4°C. For DNA staining, cell sample were incubated with SYTO9DNA fluorescent dye (Thermo Fischer Scientific) at 5 μg/mL final concentration solution and incubated in the dark for 15 minutes before analyze. Flow cytometry analysis were performed on an Attune® NxT Acoustic Focusing Cytometer (Thermo Fisher Scientific) at a ~120000 cells/min flow rate. Forward-Scattered (FSC) and Side-Scattered (SSC) light as SYTO9 dye fluorescence signal (FL-1) were acquired for at least 100,000 cells with appropriate setting (Voltage FSC 300, SSC 460 and FL1 300, threshold of FCS 500 and SSC 5000).

